# Quinone reductase 2 dimerization is dynamically driven by ligand binding

**DOI:** 10.64898/2026.05.10.723996

**Authors:** Maya Miller, Dan Loewenthal, Philipp Kukura, Nathaniel Gould

## Abstract

Human quinone reductase 2 (QR2, NQO2) is a cytosolic flavoprotein involved in cell physiology and metabolism, and implicated in several diseases. However, the mechanisms that govern its oligomeric assembly and diverse functional outcomes remain incompletely understood. Here, we employ native mass spectrometry to directly resolve the dynamic oligomeric landscape of recombinant human QR2 expressed in *Escherichia coli*, preserving non-covalent interactions and enabling analysis of assembly behavior under native conditions. QR2 is predominantly observed as a dimer stabilized by multiple non-covalently bound ligands, giving rise to discrete species. Top-down native mass spectrometry reveals a single intact proteoform, excluding covalent modification or covalently bound flavins as drivers of oligomerization. Binding of flavin adenine dinucleotide (FAD) robustly stabilizes the dimer, while unexpectedly, flavin mononucleotide (FMN) also promotes dimer formation. As FMN and FAD differ structurally by the presence of an adenine dinucleotide moiety, we hypothesized that purine nucleotide binding itself may modulate QR2 assembly. Consistent with this, we identify a new concentration-dependent effect of guanosine-triphosphate (GTP) on QR2 dimerization. Functional reductase assays show that flavin-stabilized dimers exhibit the highest catalytic activity, whereas GTP-induced dimers retain reduced activity. Binding of the inhibitor YB537 abolishes activity despite promoting dimer formation. Together, these findings reveal a ligand-dependent structural plasticity in QR2 oligomerization that is decoupled from reductase function, suggesting that QR2 dimerization serves a wider regulatory role beyond simply supporting reductase catalysis.

## 1. Introduction

Quinone reductases play a central role in cellular redox homeostasis by catalyzing the reduction of quinones and related substrates, thus limiting the generation of reactive oxygen species and contributing to detoxification pathways [1]. In humans, quinone reductase 2 (QR2), also known as NQO2, is a cytosolic flavoprotein that shares significant sequence and structural similarity with quinone reductase 1 (QR1/NQO1), but exhibits distinct biochemical properties and biological functions [2, 3]. Although QR1 is well characterized as a NAD(P)H-dependent detoxifying enzyme with broadly defined cytoprotective functions, QR2 has evolved away from the use of NAD(P)H and is involved in various context-dependent functions, including metabolic processes and neuronal physiology, among others [4]. Importantly, QR2 is also involved in toxification processes, and its dysregulation is observed in cancer and neurodegenerative diseases [5, 6]. To wit, QR2 inhibitors are neuroprotective and have rescued mice models of Alzheimer’s disease [7, 8]. Despite growing evidence linking QR2 to important physiological processes and disease contexts, the molecular mechanisms underlying QR2 regulation and function are incompletely elucidated [5].

Crystallographic and biochemical studies have shown that QR2 is a homodimeric enzyme, dependent upon flavin adenine dinucleotides (FAD) association for stability and catalytic competence [3]. Nevertheless, most of the existing structural insights into QR2 are derived from static, high-concentration conditions, providing limited information about the dynamic nature of its oligomeric assembly in solution. Consequently, it remains unclear whether QR2 dimerization is intrinsically driven by protein concentration or is instead actively regulated by small molecules. This question is especially pertinent given the many documented interactions QR2 has with a variety of endogenous and exogenous ligands, including monoamines, acetaminophen, and various naturally occurring and pharmacologically derived inhibitors [2, 5, 9–13]. However, the impact of these interactions on its quaternary structure have not been systematically explored.

Despite substantial sequence similarity between QR1 and QR2, accumulating evidence indicates that these enzymes are regulated through distinct structural and ligand-dependent mechanisms. Mechanism-based inhibition studies have demonstrated that small molecules can selectively target QR2 over QR1 by exploiting differences in the architecture of the active-site and the position of FAD, highlighting the fundamental structural divergence between the two enzymes despite their partial homology [2, 8, 14, 15]. In QR1, ligand binding has been shown to modulate enzymatic activity through conformational communication across the dimer interface, establishing that functional output depends not only on dimerization but also on the specific conformation adopted by the dimeric assembly [16]. These observations provide a conceptual framework for understanding how ligand-induced remodeling of the QR2 quaternary structure may regulate reductase function.

Native mass spectrometry (nMS) provides a powerful tool to interrogate protein assembly and ligand binding under physiological conditions [17, 18]. By preserving non-covalent interactions during ionization, nMS enables direct observation of coexisting oligomeric and ligand-bound species, allowing subtle shifts in equilibrium to be resolved without the need for labeling or immobilization [18, 19]. As such, nMS is uniquely suited to dissecting ligand-regulated assembly mechanisms in proteins whose functional states may depend on dynamic oligomerization.

In this study, we applied nMS to investigate the oligomerization behavior of recombinant human QR2 expressed and purified from *Escherichia coli*. We examine how flavin cofactors, nucleotides, and a selective small-molecule inhibitor influence monomer-dimer formation. By integrating nMS with a top-down approach and functional enzymatic assays, we demonstrate that QR2 assembly is largely independent of protein concentration and instead is tightly regulated by non-covalent ligand binding. This represents a central regulatory mechanism that links the function of QR2 with the cellular chemical environment, particularly in the context of brain physiology and disease.

## 2. Results and Discussion

### 2.1. Ligand-associated oligomerization of human QR2 revealed by native mass spectrometry

Recombinant human QR2 was heterologously expressed in *Escherichia coli*, which provides a robust expression platform that shares significant overlap with small molecules found in mammalian cells [20]. QR2 was purified by nickel-affinity chromatography without further sample manipulation or deliberate cofactor supplementation. nMS of the purified protein revealed two well-resolved charge state distributions corresponding to monomeric and dimeric QR2, consistent with their respective molecular masses (Figure 1A,B). In addition to apo forms, the dimeric population exhibited pronounced heterogeneity, characterized by multiple discrete mass shifts indicative of small molecule binding.

**Figure 1.**
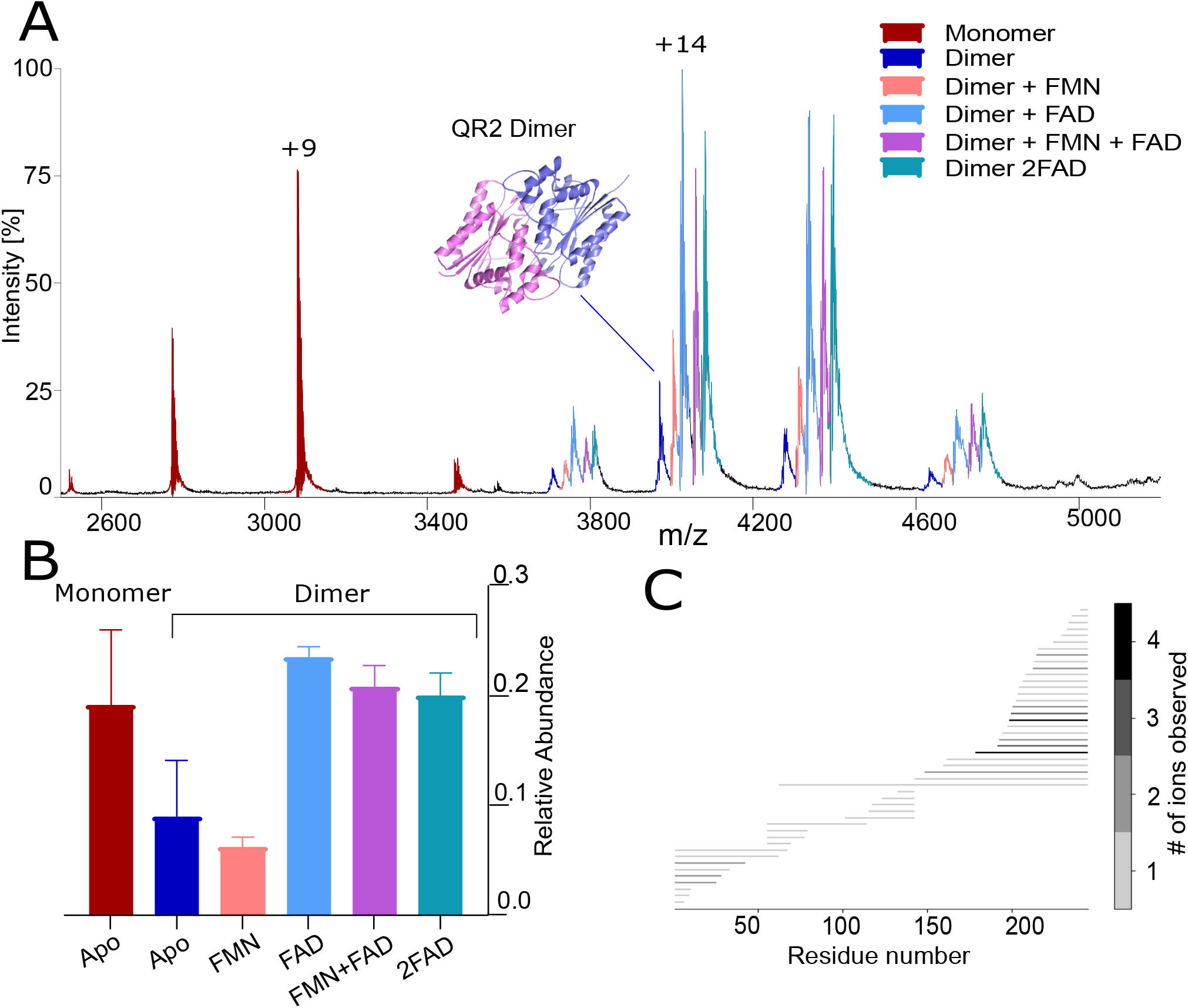
Native mass spectrometry characterization of QR2 and associated ligand-binding states. **(A)** Native mass spectrum of recombinant human QR2 showing coexisting monomeric (red) and dimeric species (color-coded by ligand-binding state). The dimeric region reveals apo dimer (dark blue) and four predominant non-covalent adducts corresponding to binding of a single FMN (salmon), a single FAD (light blue), a mixed FMN-FAD species (purple), and two FAD molecules (cyan) per QR2 dimer. An inset shows the predicted QR2 homodimer structure generated using AlphaFold [22]. **(B)** Relative abundance of the individual dimer-associated binding states derived from native mass spectra. **(C)** Top-down native mass spectrometry of intact QR2 showing fragment ion coverage across the protein sequence, indicating the absence of covalently bound small molecules or post-translational modifications.

Four distinct dimer-associated adducts were reproducibly observed, corresponding to mass increases of +456, +786, +1241 and +1570 Da relative to the apo dimer. Based on mass agreement and known cofactor masses, the lowest-mass adduct is consistent with binding of FMN, while the +786 Da adduct corresponds to a single FAD molecule. The +1241 Da species are consistent with the simultaneous binding of FAD and FMN, while the highest-mass adduct corresponds to the occupancy of two FAD molecules. The presence of these multiple, well resolved dimeric species in the absence of exogenous ligand addition indicates that QR2 purified from *Escherichia coli* exists as an ensemble of cofactor-associated states and suggests that various endogenous small molecules can associate and stabilize different dimeric assemblies.

The native mass spectrum shows that QR2 is predominantly observed in the dimeric assembly, indicating that the dimer represents the principal oligomeric form of the protein in solution (Figure 1B). Quantitative analysis of the dimer-associated species revealed that only a minor fraction corresponds to the apo dimer, whereas flavin-associated dimers dominate the spectra. In particular, FAD-containing dimers constitute the most abundant population, appearing either as single FAD-bound dimers, mixed FAD-FMN species, or dimers occupied by two FAD molecules. These observations indicate that flavin binding strongly stabilizes the QR2 dimer and suggest that the surrounding small-molecule environment can shape the composition and relative abundance of ligand-associated dimeric states.

To determine whether covalent proteoform heterogeneity contributes to this behavior, intact QR2 was further analyzed by top-down nMS (Figure 1C and Supplementary Figure S1). Spectra acquired on an Orbitrap mass spectrometer using higher-energy collisional dissociation (HCD) revealed a single dominant intact proteoform. Deconvolution using PrecisION [21] yielded a molecular mass consistent with the theoretical sequence mass within ±3 Da. Fragment ion analysis provided 18% sequence coverage, with ions distributed throughout the N-terminus, internal regions, and C-terminus, as confirmed by fragment position and coverage maps (See Supplementary Figure S1). No mass shifts or fragmentation patterns consistent with post-translational modifications or covalently bound cofactors were detected at any position along the sequence. These data exclude covalent modification, including rare covalent flavin attachment, as contributors to QR2 oligomerization.

Collectively, these results demonstrate that recombinant human QR2 exists as a heterogeneous ensemble of non-covalently ligand-associated oligomeric states. The presence of multiple cofactor-bound dimer species, together with the absence of covalent modification, indicates that QR2 assembly is likely to be governed by its chemical microenvironment.

### 2.2. Structural analysis of ligand interactions with QR2

To investigate how ligand binding contributes to QR2 dimerization, we first analyzed flavins associated with the predominant dimeric species. Comparison of the chemical structures of FAD and FMN (Figure 2A) shows that FAD comprises an FMN core attached to an adenine nucleotide moiety. This raises the possibility that purine nucleotide-like interactions absent in FMN but present in FAD could contribute to dimer stabilization and motivate the interrogation of purine nucleotides themselves as potentially novel modulators of QR2 dimer assembly.

**Figure 2.**
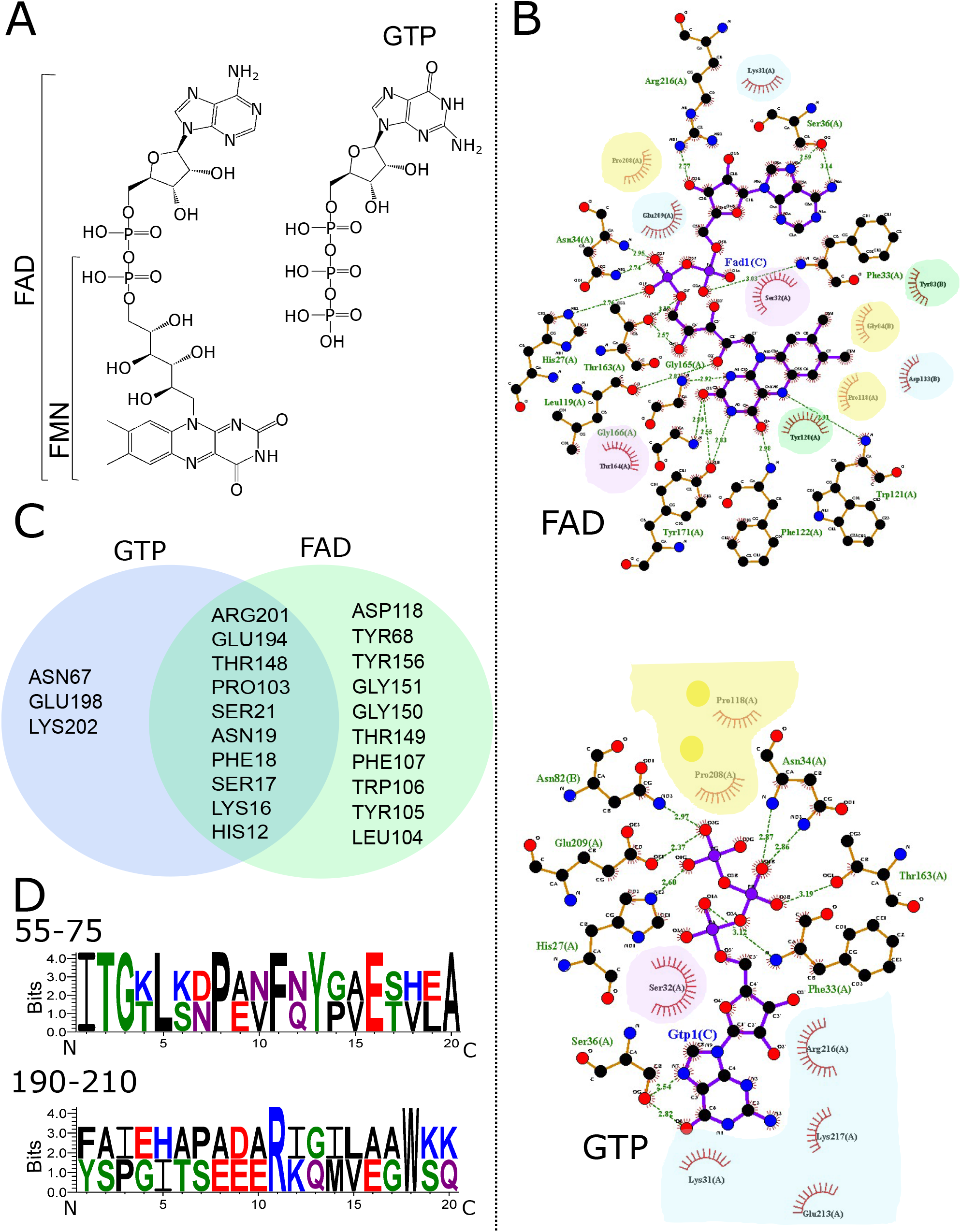
Structural comparison of FAD and GTP interactions with QR2. **(A)** Chemical structures of FMN, FAD, and GTP. **(B)** LigPlot [23] analyses showing QR2 residues predicted to interact with GTP and FAD based on AlphaFold [22] structural models. Hydrogen bonds and non-bonded contacts are indicated. Residues are color-coded according to chemical properties: blue denotes electrically charged residues, green hydrophobic residues, purple uncharged polar residues, and yellow conformationally distinctive residues (Gly, Pro, Cys). **(C)** Venn diagram summarizing predicted QR2 residues involved in FAD and GTP binding, highlighting shared and ligand-specific interaction sites. **(D)** Sequence alignment of QR1 (top) and QR2 (bottom) across two 20-residue windows (positions 55-75 and 190-210) located proximal to the predicted ligand-binding region created with WebLogo [24].

Therefore, we analyzed the QR2 interactions predicted by Alphafold [22] with FAD, ATP and GTP using LigPlot [23]. This revealed that while there is significant overlap between the interactions these ligands have with QR2 (Supplementary Figure S2), GTP and FAD engage QR2 through chemically distinct interaction networks (Figure 2B). GTP binding is dominated by electrostatic contacts with charged residues (Arg216, Lys217, Glu213, Lys31) and the involvement of conformationally distinctive residues (Pro208, Pro118), consistent with stabilization of nucleotide triphosphate through charge complementarity and local backbone flexibility. In contrast, FAD engages in a mixed environment comprising polar, aromatic and structural residues (Thr164, Tyr120, Tyr83, Gly84, Asp133, Ser32, Glu209, Pro208, Lys31), characteristic of canonical flavoprotein binding pockets. Venn analysis identified 11 shared residues (Figure 2C), in addition to 10 FAD-specific and only 3 GTP-specific contacts, indicating partial spatial overlap, but fundamentally different binding chemistries. The limited number of unique GTP residues suggests a comparatively permissive binding mode, whereas the broader FAD-specific interaction network is consistent with a more rigid and structurally defined binding mechanism. Next, we compared the QR1 and QR2 sequences in two 20-residue windows proximal to the predicted ligand-binding region to assess the extent of local conservation between the proteins (Figure 2D). Although several residues are retained, both regions differ in their local sequence composition and chemical character. In particular, the 55-75 region exhibits partial similarity between QR1 and QR2, while the 190-210 region is predominantly divergent. Interestingly, this latter region contains two of the three residues identified as GTP-specific in the LigPlot analysis, directly linking local sequence divergence to the nucleotide-binding interface. These observations suggest that sequence variation in this region underpins the distinct GTP-binding mode and contributes to the specific ligand remodeling of the QR2 dimer. More broadly, these substitutions are likely to reshape the ligand-proximal environment, providing a structural basis for the pronounced ligand-dependent plasticity of QR2 despite the conservation of the overall fold and dimer architecture.

Collectively, these findings support a model in which chemically distinct ligands engage overlapping but non-identical residue networks to stabilize multiple, functionally distinct dimeric states. In this framework, the availability of ligands emerges as a key determinant of QR2 oligomerization, offering a mechanistic basis for how cofactors, nucleotides, and inhibitors may differentially remodel QR2 structure and function.

### 2.3. Ligand-dependent remodeling of QR2 dimer assembly

Based on our *in silico* analysis and nMS findings, we tested the effects of nucleotide (GTP) and flavins (FAD and FMN) on QR2 assembly by titrating ligands and monitoring changes in dimer formation using nMS. The addition of GTP induced a shift towards the formation of QR2 dimers (Figure 3A,B), revealing a previously unreported nucleotide sensitivity of QR2 dimer assembly. Rather than collapsing the population into a single holo-dimer, GTP promoted redistribution among existing dimer-associated species, consistent with a graded and tunable stabilization mechanism. The observed mass increment is formally consistent with the binding of two FAD molecules or three GTP molecules. However, since no exogenous FAD was supplied, the GTP-dependent redistribution strongly suggests displacement of pre-associated FAD by GTP. Although a three-GTP stoichiometry is unexpected, the observation of odd-numbered endogenous adducts in the native spectra suggests that QR2 can accommodate asymmetric and non-integer cofactor occupancies, making higher-order nucleotide binding plausible. This heterogeneity points to a structurally modular dimer capable of engaging multiple small molecules in noncanonical arrangements, indicating that the assembly of QR2 involves more complex ligand coordination than simple symmetric ligand/cofactor binding. Importantly, GTP binding caused a shift in the m/z distribution of the dimer relative to native QR2 in the absence of a net mass change, which may indicate ligand-induced conformational rearrangement of the QR2 dimer rather than simple exchange of cofactors.

**Figure 3.**
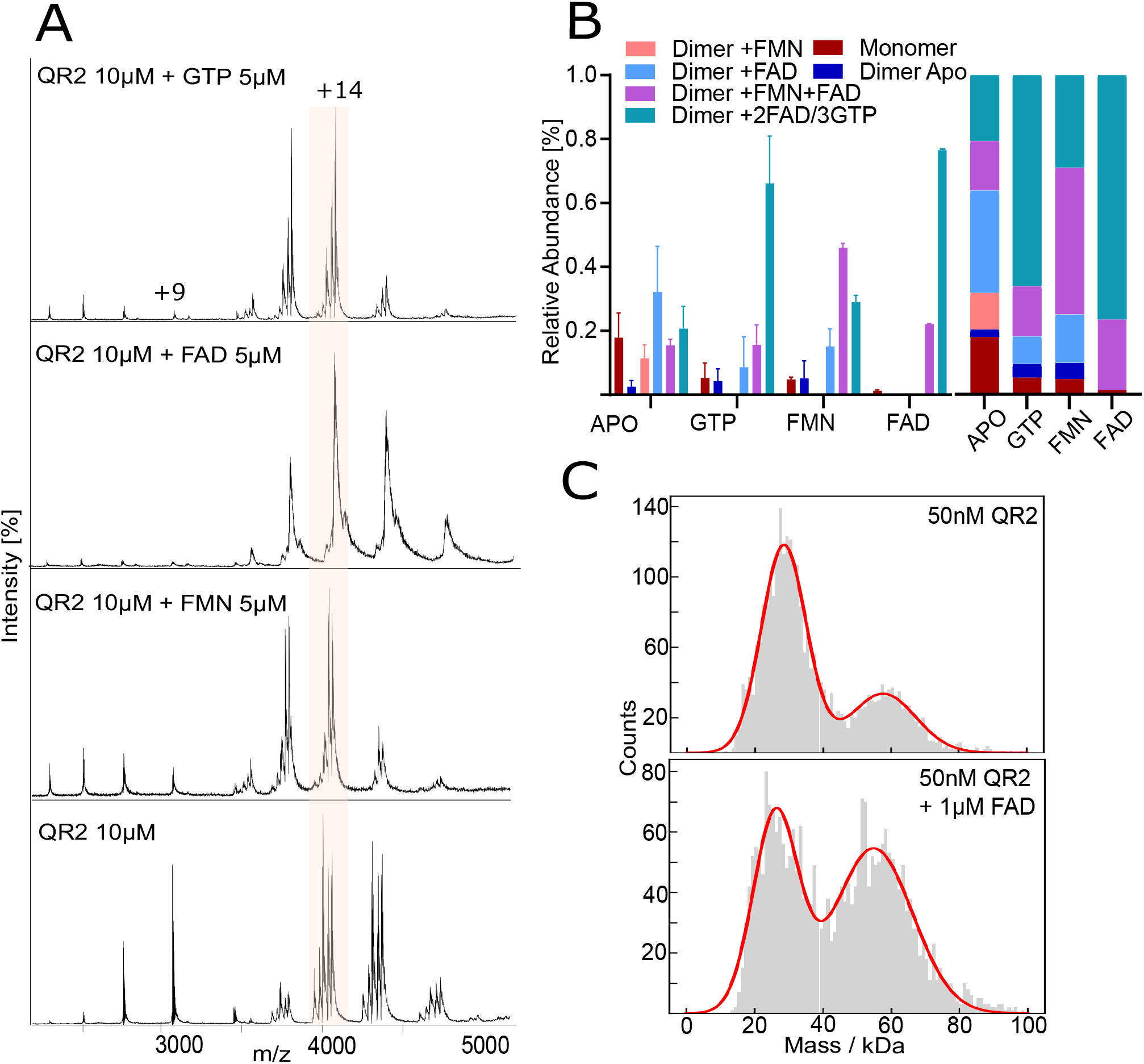
Ligand-dependent remodeling of QR2 oligomeric states measured by native mass spectrometry and mass photometry. **(A)** Native mass spectra of QR2 (10 µM) following incubation with 5 µM of GTP, FAD, and FMN. Each ligand induces redistribution of QR2 dimer-associated species relative to the untreated protein. **(B)** Quantification of relative abundances of monomeric and dimeric QR2 species under each condition. Bar plots show mean ± SD across replicates presented as relative abundance and as stacked bar charts to illustrate redistribution. **(C)** Mass photometry measurements of QR2 (50 nM) showing the oligomeric distribution under apo conditions (upper panel) and following addition of 1 *μ*M FAD (lower panel). The monomeric and dimeric populations correspond to signals centered at approximately 28 kDa and 56 kDa, respectively.

Consistent with previous biochemical and structural studies, the addition of FAD produced a pronounced shift toward a dominant holo-dimeric species (Figure 3A,B), confirming its role as a primary stabilizing prosthetic group. The binding of FAD effectively displaced alternative dimeric populations, yielding a highly uniform assembly and serving as an internal validation of the nMS approach. In contrast, FMN induced a markedly different response (Figure 3A,B and Supplementary Figure S3). FMN promoted dimer formation but simultaneously drove extensive remodeling of cofactor occupancy, characterized by enrichment of FMN-associated dimers and reciprocal depletion of FAD-bound species. This competitive exchange demonstrates that FMN can partially replace pre-associated FAD, generating a heterogeneous ensemble of dynamically interconverting dimeric states. Unlike FAD, which enforces a rigid holo-dimer, FMN engages QR2 more transiently, fine-tuning assembly rather than locking it into a single configuration. This reveals previously unknown functional aspects in the relationship between QR2 and FMN.

To independently validate the oligomeric behavior observed by nMS, QR2 was analyzed using mass photometry, an orthogonal technique that directly reports protein stoichiometry in solution [25, 26]. Measurements performed on QR2 revealed a mixture of monomeric and dimeric species under apo conditions. Upon the addition of FAD, a pronounced shift was observed toward the dimeric population, accompanied by the depletion of the monomeric species (Figure 3C). These results closely mirror the enrichment of ligand-dependent dimers detected by nMS and confirm that flavin binding stabilizes QR2 dimers in solution. These combined findings show that ligand binding promotes QR2 dimerization under physiological conditions.

Overall, these results reveal a hierarchy of ligands in the QR2 assembly, in which GTP introduces nucleotide-sensitive modulation, FAD imposes strong structural stabilization, and FMN enables dynamic cofactor exchange. In particular, FMN and GTP binding induce reproducible shifts in the QR2 dimer m/z envelope without corresponding changes in intact mass, consistent with ligand-driven conformational remodeling that alters charge-state distribution. In contrast, FAD binding produces a single dominant holo-dimer species without detectable m/z redistribution, indicative of a more rigid, structurally constrained assembly. This spectrum of responses highlights the pronounced structural heterogeneity in QR2 dimerization, demonstrating that chemically distinct ligands remodel QR2 into multiple coexisting dimeric conformers. Collectively, these findings establish QR2 as a ligand-responsive assembly whose quaternary structure is tuned by identity and availability of ligands, rather than protein concentration alone, revealing conformational heterogeneity as an integral characteristic of QR2 regulation.

### 2.4. Ligand-dependent QR2 dimerization uncouples oligomeric state from catalytic activity

To directly relate ligand-induced QR2 dimerization to enzymatic function, we compared oligomeric state and reductase activity in the presence of FMN, FAD, GTP, and QR2 inhibitor YB537 [8] under matched conditions. First, nMS of YB537 with QR2 showed that, similarly to flavins and GTP, incubation with YB537 resulted in the formation of QR2 dimers (Figure 4A,B and Supplementary Figure S4). However, in contrast to the distinct distribution of dimer-associated peaks and characteristic shifts along the m/z axis described for GTP and flavins, YB537 caused enrichment of higher-mass dimer species consistent with binding of three inhibitor molecules per QR2 dimer. This differs from the two FAD molecules observed under flavin-saturated or non-flavin-saturated conditions. This altered stoichiometry further supports ligand-dependent remodeling of the QR2 dimer and suggests that YB537 engages QR2 through a binding mode distinct from that of flavin association, and provides new details regarding YB537 binding to QR2, beyond those revealed by the resolved crystal [8].

**Figure 4.**
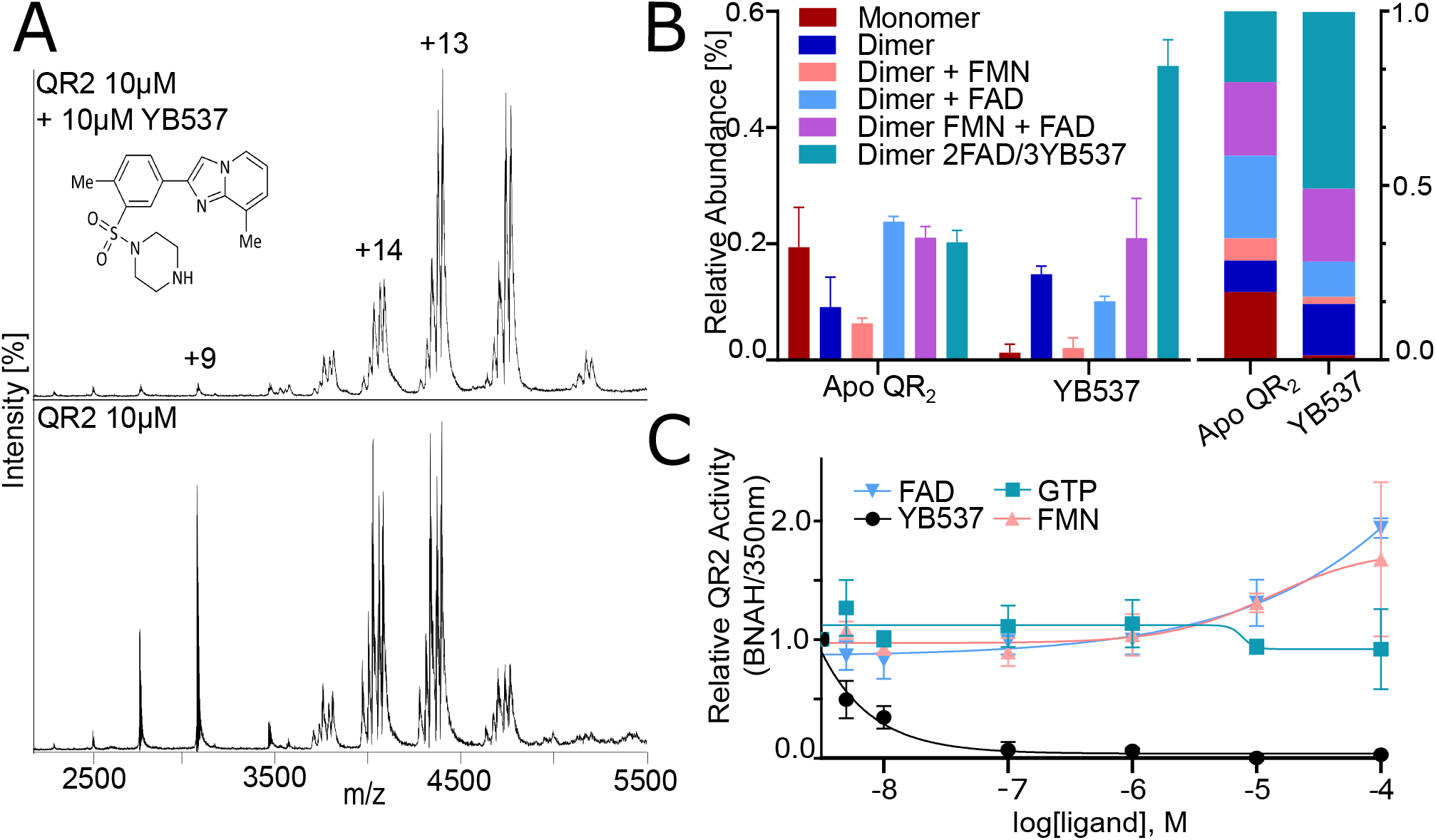
Modulation of QR2 dimer composition and reductase activity by YB537 and small-molecule ligands. **(A)** Native mass spectra of QR2 (10 µM) following incubation with YB537 (10 µM) and chemical structure of the QR2 inhibitor YB537. **(B)** Quantitative analysis of dimer-associated species, presented as relative abundance and stacked bar charts to illustrate redistribution of cofactor-bound dimer populations upon inhibitor binding. **(C)** Functional reductase assays of QR2 in the presence of GTP, YB537, FAD, and FMN, showing ligand-dependent modulation of catalytic activity.

To assess whether these structurally distinct dimeric assemblies exhibit corresponding differences in catalytic competence, QR2 activity was measured using a spectrophotometric assay driven by the artificial hydride donor N-benzyldihydronicotinamide (BNAH)- and acceptor, menadione, and the reaction progress was monitored at 350 nm (Figure 4C) [8]. In the presence of flavin cofactors, both FAD- and FMN-bound QR2 displayed robust enzymatic activity. Although FAD produced the most homogeneous dimeric population as measured by nMS, FMN-supported dimers retained comparable catalytic output, indicating that variations in cofactor occupancy and dimer composition do not preclude reductase activity.

In contrast, the addition of GTP produced an attenuation of QR2 activity relative to flavin-bound states, consistent with nMS observations showing redistribution of cofactor-associated dimer species rather than complete displacement of catalytically competent flavins. As expected, YB537 prevented reductase activity, despite strongly stabilizing the dimeric assembly. This dissociation between dimer formation and catalytic output demonstrates that not all QR2 dimers are functionally equivalent.

Together, these results establish that QR2 dimerization alone is not predictive of reductase activity. Instead, ligand identity dictates both the structural properties of the dimeric assembly and its functional competence. Flavins promote catalytically active dimers, GTP induces dimerization with reduced activity, and YB537 binding stabilizes inactive dimeric states. The observation that chemically distinct ligands universally enrich dimer formation but produce markedly different enzymatic profiles supports a model in which QR2 exists as a spectrum of ligand-defined dimer conformations, with functional output determined by small-molecule engagement rather than oligomeric state per se.

## 3. Discussion and Conclusion

In this study, we show that human QR2 exists not as a single static holoenzyme, but as a ligand-responsive assembly whose quaternary structure is dynamically shaped by small-molecule engagement. Using nMS, we resolve multiple coexisting QR2 dimeric states stabilized by flavins, nucleotides, and the QR2 inhibitor YB537, revealing a previously unrecognized degree of structural plasticity in this enzyme. Rather than being governed solely by protein concentration, QR2 assembly is primarily dictated by ligand availability, highlighting small molecules as dominant regulators of QR2 architecture.

Our data demonstrate that stable QR2 dimerization can occur upon binding of a single flavin cofactor, indicating asymmetric occupancy and intrinsic flexibility at the dimer interface. This contrasts with classical models of flavoprotein assembly that assume symmetric holo-dimer formation and suggests that QR2 can adopt partially occupied, structurally heterogeneous states. This behavior is consistent with a growing appreciation that many metabolic enzymes populate ensembles of ligand-defined conformations rather than discrete on-off states [27].

Beyond flavins, we identify GTP as a previously unreported modulator of QR2 assembly. GTP induces dimerization while simultaneously remodeling the cofactor-bound population and shifting the m/z distribution without a corresponding change in intact mass, consistent with ligand-driven conformational rearrangement of the QR2 dimer. Although the precise nucleotide stoichiometry remains ambiguous, the observed redistribution strongly suggests displacement of endogenous flavins by GTP and highlights nucleotide binding as an additional regulatory axis for QR2. These findings expand the functional landscape of QR2 beyond redox cofactors and implicate the availability of cellular nucleotides as a potential determinant of the structural state of QR2.

Comparative *in silico* analysis further reveals that FAD and GTP engage QR2 through chemically distinct interaction networks. Whereas FAD binding involves a broad constellation of residues characteristic of canonical flavoprotein pockets, GTP interacts through a more limited and predominantly charged residue set, indicative of a comparatively permissive binding mode. This difference in binding chemistry provides a structural rationale for our nMS observations that FAD enforces a rigid, homogeneous holo-dimer, while GTP promotes a more plastic assembly capable of accommodating multiple ligand configurations. Together with the dynamic exchange observed for FMN, these results support a model in which QR2 integrates chemically diverse ligands into overlapping but non-identical binding environments to access multiple quaternary states.

Importantly, ligand-induced dimerization is uncoupled from catalytic competence. Although flavins, GTP, and YB537 all stabilize dimeric QR2, only flavin-bound assemblies support robust reductase activity, whereas GTP partially attenuates activity and inhibitor binding abolishes it entirely. These findings establish that dimer formation alone is not predictive of enzymatic function and instead that ligand identity defines the functional output of the QR2 dimer. This uncoupling suggests that QR2 dimers may serve broader regulatory or signalling roles beyond supporting quinone reduction, potentially enabling QR2 to operate as a ligand-responsive scaffold or molecular sensor [4, 11, 28]. Evidence for this is growing, and a recent study demonstrates a function for QR2 in epigenetic regulation, by being a reader of histone H3 serotonylation [28].

These observations also align with emerging evolutionary and biochemical evidence that QR2 differs fundamentally from its paralog QR1 [4], including markedly reduced catalytic efficiency with physiological nicotinamide cofactors and evolutionary selective pressure toward diminished redox activity. Together, these findings raise the possibility that QR2 may function in cells not solely as a reductase but as a structurally adaptive protein whose biological roles are mediated through ligand-dependent assembly states. Such a framework may help reconcile QR2’s reported involvement in neurological processes, metabolic regulation, and disease phenotypes, where signaling and protein-protein interactions rather than redox chemistry may predominate. Future studies using human cells, to analyze QR2 interactions by nMS in different cellular compartments in physiological and disease contexts, would be needed to fully explore these possibilities.

More broadly, our work illustrates the power of nMS to resolve dynamic protein assemblies and uncover regulatory mechanisms that remain inaccessible to static structural methods. By preserving non-covalent interactions and directly reporting on ligand occupancy and oligomeric state, nMS enables dissection of assembly landscapes that underpin functional diversity.

In summary, we propose that QR2 exists as a spectrum of ligand-defined dimeric conformations whose structural and functional properties are tuned by the chemical environment. This ligand-dependent plasticity provides a mechanistic basis for QR2 regulation and suggests new strategies for therapeutic intervention, in which targeting assembly equilibria rather than active sites alone may offer more effective modulation of QR2 biology.

## Materials and Methods

### Protein expression and purification

The plasmid encoding human quinone reductase 2 (hNQO2; NM000904.6) with an N-terminal 6×His tag followed by a TEV protease cleavage site was obtained in the bacterial expression vector pET-6xHis / TEV (Vector ID: VB240530-1459jna). Plasmids were amplified by transforming into *E. Coli* UltraStable cells and confirmed by Sanger sequencing. For protein expression, the verified plasmid was transformed into chemically competent *E. coli* BL21(DE3) cells. Overnight preculture was grown at 37 °C by inoculating a freshly transformed colony in a 100 ml LB media. The preculture was diluted 1:100 into fresh 6 × 1 L LB media and was grown at 37 °C until the OD600 reached 0.6. Cells were induced by the addition of 0.5 mM isopropyl *β*-D-1-thiogalactopyranoside (IPTG), and the culture was grown for another 3 hours at 37 °C. Cells were pelleted using centrifugation with the Beckman JLA 8.1000 rotor for 15 minutes at 5000 × g. The pellet was resuspended in lysis buffer consisting of 20 mM Tris-HCl (pH 8.0), 150 mM NaCl, and 10% (v/v) glycerol, supplemented with an EDTA-free protease inhibitor cocktail. The lysed cells were then centrifuged at 4 °C to separate the insoluble material using a JA-25.50 rotor at 20,000 × g, 30 min. The clarified supernatant was collected and subjected to His-tag affinity purification to isolate recombinant hNQO2.

### Native mass spectrometry and top-down mass spectrometry

Purified QR2 was buffer-exchanged into 200 mM ammonium acetate (pH 8.0) using gel-filtration spin columns (75 *μ*L Zeba Spin Desalting Columns; Thermo Fisher Scientific) immediately prior to analysis. For ligand-spiking experiments, YB537 [8], GTP (Sigma-Aldrich, G8877), FAD (Fisher Scientific, 11411838), and FMN (Cambridge Bioscience, 332-10509-2) were individually weighed and dissolved directly in 200 mM ammonium acetate (pH 8.0) to prepare 1 mM stock solutions. Ligands were then diluted to the indicated final concentrations directly into the QR2 samples immediately prior to native mass spectrometric analysis, as specified in the corresponding spectra.

Native mass spectrometry measurements were performed on a Q Exactive UHMR mass spectrometer (Thermo Fisher Scientific) operated in positive ion mode using the manufacturer’s recommended settings for native mass spectrometry. Spectra were acquired at a resolving power of 12,500 (at *m*/*z* 200), with HCD set to 10, in-source trapping (IST) set to 50, and an acquisition range of *m*/*z* 350–10,000. Nanoelectrospray ionization was achieved using in-house prepared gold-coated borosilicate capillaries supplied with a slight backing pressure (approximately 0.5 mbar) and held at 1.0 kV relative to the instrument inlet. The inlet capillary temperature was maintained at 200 °C.

Top-down mass spectrometry was performed on intact QR2 using an Orbitrap Eclipse Tribrid mass spectrometer (Thermo Fisher Scientific) operated in positive ion mode with higher-energy collisional dissociation (HCD). Native protein ions were generated by nanoelectrospray ionization using a spray voltage of 1.0 kV relative to the instrument sampling interface (200 °C), with 200 V in-source activation applied for desolvation. The instrument was operated in intact protein mode under high-pressure conditions. MS1 spectra were acquired over an *m*/*z* range of 500–8,000 at a resolving power of 120,000. MS2 spectra were acquired using HCD with a collision energy of 200 V.

Raw spectra were processed and deconvoluted using PrecisION software [21]. Observed intact masses were compared with the theoretical sequence mass to assess proteoform heterogeneity using a mass tolerance threshold of ±3 Da.

### Enzymatic Activity

The activity of QR2 reductase was quantified by measuring the oxidation rate of the synthetic cofactor benzyl dihydro-nicotinamide (BNAH), as previously described [8]. Briefly, a final concentration of 100*μ*M BNAH (Fluorochem, F545103) and 100*μ*M menadione (Fluorochem, F049845) were dissolved into a buffer of 50mM Tris with 150mM NaCl at pH 8.5, in the presence of 10nM recombinant human QR2, with increasing concentrations of YB537 [8], GTP (Sigma-Aldrich, G8877), FAD (Fisher Scientific, 11411838), or FMN (Cambridge Bioscience, 332-10509-2). BNAH oxidation was measured by absorption at 350nm in a reaction volume of 120*μ*L with a PHERAstar FSX plate reader (BMG Labtech), using a 96 well clear plate format (Scientific Laboratory Supplies, Corning-3997). The initial velocity was determined and compared across conditions. The activity was measured in triplicates.

### Mass Photometry

Measurements were performed on a TwoMP mass photometer (Refeyn). Glass coverslips (Epredia no. 1.5 coverslips, 50 × 24 mm, VWR, cat. no. 16002-264) were cleaned by 5 min ultrasonication in 50:50 IPA:MQ (24137-2.5L-M, Sigma Aldrich), followed by 5m ultrasonication in Milli-Q (MQ) water. Silicone gaskets (CultureWell™, CW-50R-1.0, 50–3 mm diameter × 1 mm depth) were placed on the coverslips. Samples were measured in PBS at a final concentration of 50 nM QR2, in the presence or absence of 1 *μ*M FAD. Data were acquired for 60 s at 237 frames s^−1^ using AcquireMP (Refeyn, 2023 R1.1). The resulting movies were analysed using DiscoverMP (Refeyn, v2024 R1).

## Author Information

### Authors

Dan Loewenthal — Department of Chemistry, University of Oxford, OX1 3TA, UK; Kavli Institute for Nanoscience Discovery, University of Oxford, OX1 3QU, UK

Philipp Kukura — Department of Chemistry, University of Oxford, OX1 3TA, UK; Kavli Institute for Nanoscience Discovery, University of Oxford, OX1 3QU, UK

### Author Contributions

M.M. and N.G. conceived the project, performed experiments, analyzed data, and wrote the manuscript. D.L. performed the mass photometry experiments and conducted the associated data analysis. N.G., P.K., and M.M. supervised the research and contributed to interpretation of the data. All authors discussed the results and contributed to the final manuscript.

### Funding

N.G. acknowledges support from the University of Oxford Medical Sciences Division Bridging Grant Scheme (Grant Ref: 0017765).

M.M. acknowledges support from UK Research and Innovation (UKRI) under the UKRI Postdoctoral Guarantee Scheme (Grant Ref: EP/X027236/1), following successful evaluation through the Marie Skłodowska-Curie Actions fellowship programme of the European Commission.

D.L. was supported by a Clarendon Scholarship, a Menasseh Ben Israel Scholarship, and a Kingsgate Scholarship.

P.K. acknowledges support from the Engineering and Physical Sciences Research Council (EPSRC) (EP/T03419X/1 and EP/W001055/1).

### Notes

P.K. is an academic founder, shareholder, and non-executive director of Refeyn Ltd.

## Acknowledgments

The authors thank Prof. Carol Robinson and Dr. Jani Bolla for fruitful discussions and Miss Xinye Zhang for assistance with protein production.

## Notes

### Competing Interest Statement

Philipp Kukura is an academic founder, shareholder, and non-executive director of Refeyn Ltd.

