## Supplementary for "Quinone reductase 2 dimerization is dynamically driven by ligand binding"

### Supporting Information: Quinone reductase 2 dimerization is dynamically driven by ligand binding

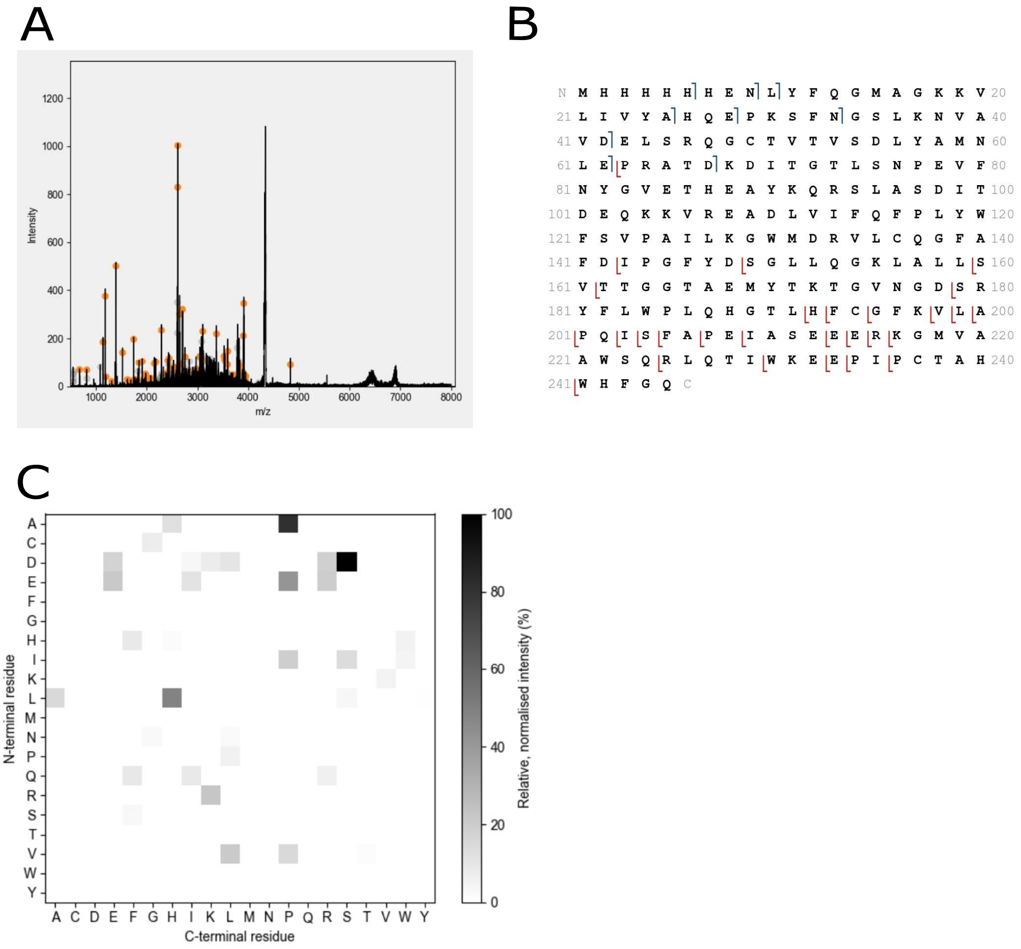


**Figure S1.** Native top-down mass spectrometry analysis of QR2. **(A)** Top-down mass spectrum chromatogram. Orange peaks represent deconvoluted ions, while gray peaks correspond to non-deconvoluted ions. **(B)** Sequence map coverage derived from top-down mass spectrometry, indicating identified fragment ions across the protein sequence. **(C)** PrecisION[1] residue-pair heatmap for UniProt P16083 (+His-tag), illustrating high-confidence residue pair assignments.

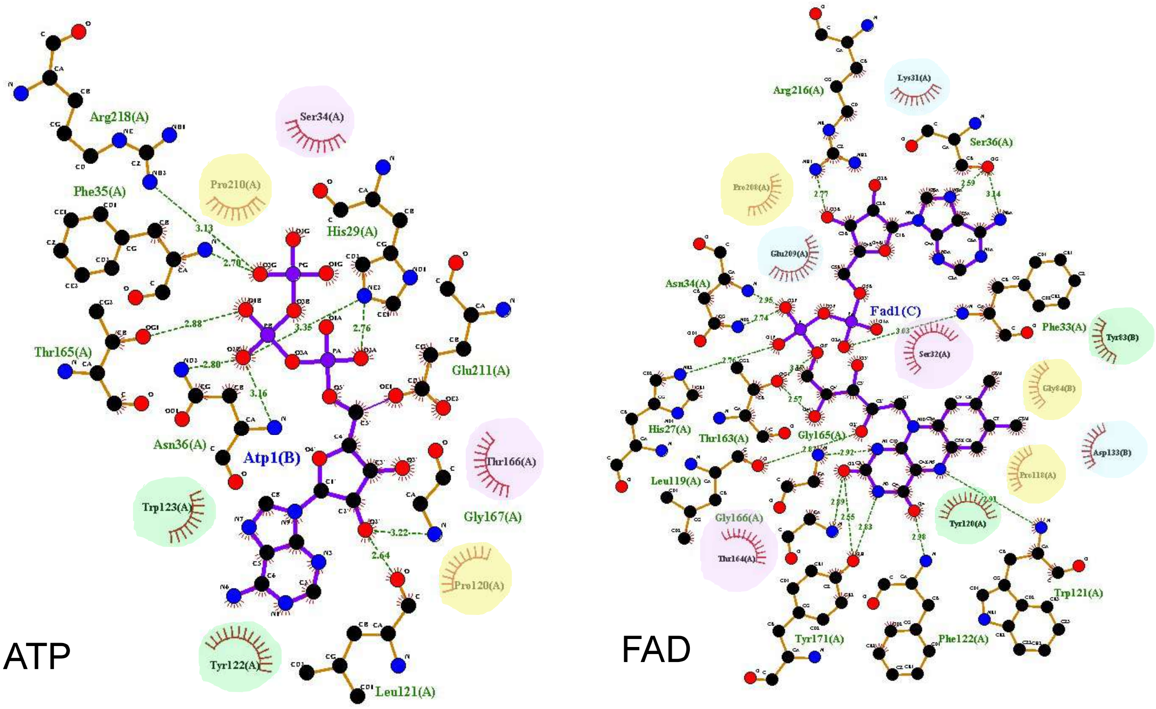


**Figure S2.** LigPlot+[2] analyses of QR2 interactions with ATP and FAD based on AlphaFold-predicted structural models.[3] Hydrogen bonds and non-bonded contacts are indicated. Residues are color-coded according to chemical properties: blue denotes electrically charged residues, green hydrophobic residues, purple uncharged polar residues, and yellow conformationally distinctive residues (Gly, Pro, Cys).

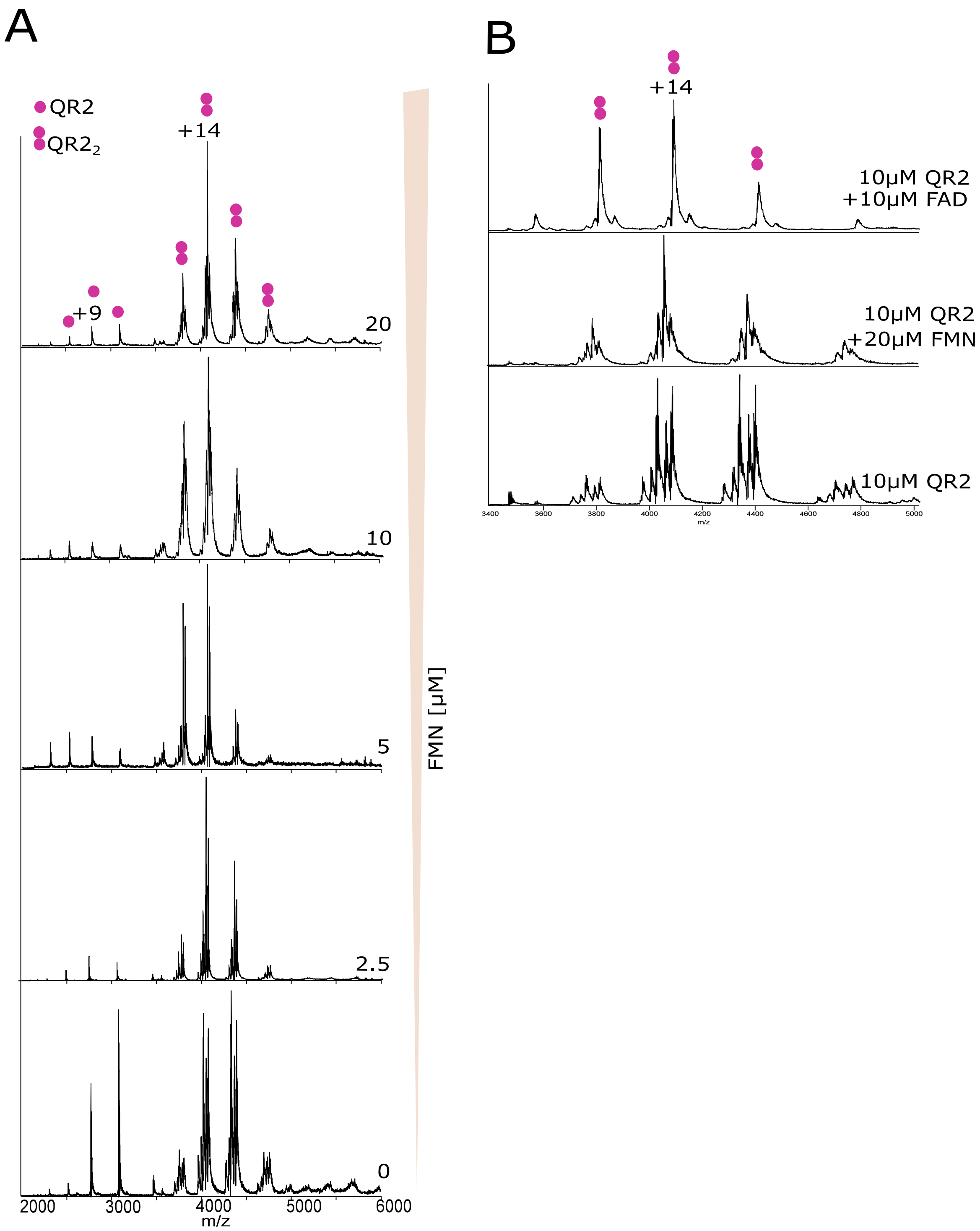


**Figure S3.** **(A)** Native mass spectrum of 10 $\mu$M QR2 spiked with increasing concentrations of FMN (0–20 $\mu$M). **(B)** Comparison of native mass spectra of QR2 alone, QR2 with 20 $\mu$M FMN, and QR2 with 10 $\mu$M FAD.

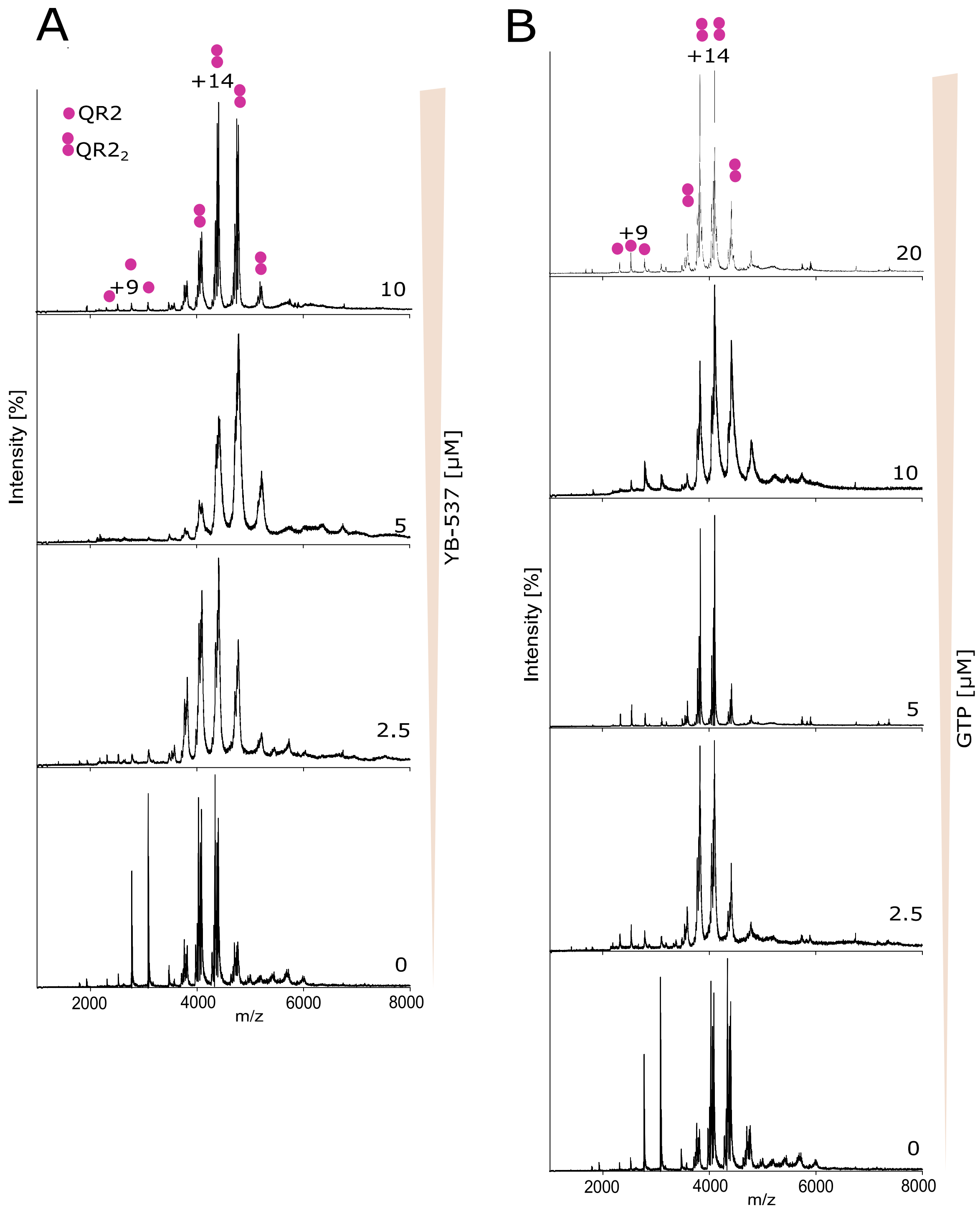


**Figure S4. (A)** Native mass spectrum of 10 $\mu$M QR2 spiked with increasing concentrations of inhibitor YB537 (0–10 $\mu$M). **(B)** Native mass spectrum of 10 $\mu$M QR2 spiked with increasing concentrations of GTP (0–20 $\mu$M).
